# LizardMorph: A generalizable machine learning framework for automated anatomical landmark detection in digital images

**DOI:** 10.64898/2026.06.10.731351

**Authors:** Mercedes Quintana, Le Yang Loh, Ayush Parikh, Jonathan J. Suh, Victoria Chávez, Arthur Porto, Breanna Shi, James T. Stroud

## Abstract

Morphological measurements underpin a wide range of ecological and evolutionary research, yet the manual landmarking workflows on which most morphometric studies depend remain a persistent bottleneck that limits both the pace and scale of biological research. Machine learning offers compelling solutions, but most automated landmarking tools require substantial computational expertise, creating a gap between technical capability and practical adoption by biologists. Here, we present LizardMorph, an integrated machine learning pipeline and web-based interface for semi-automated anatomical landmark detection on biological images. LizardMorph couples a fine-tuned ML-Morph shape predictor with an accessible, browser-based interface that enables researchers to upload images, review automated landmark predictions, interactively correct outliers through point-and-click editing, and export results in standard morphometric formats—all without programming expertise or local software installation. Using dorsal X-ray radiographs of *Anolis* lizards with 34 anatomical landmarks as a proof-of-concept, we show that the ML-Morph model achieves high predictive accuracy, with landmarks on well-defined skeletal structures predicted with 100% accuracy within a 1 mm tolerance threshold. A controlled user study comparing LizardMorph against traditional manual landmarking (TpsDig2) demonstrated significant efficiency gains: experienced annotators completed LizardMorph landmark verification 37.5% faster than manual annotation. Extrapolated to batch processing 1,000 lizards, LizardMorph saves experienced researchers approximately 6.5 hours of manual processing time. Critically, LizardMorph implements a human-in-the-loop design in which automated predictions serve as editable starting points, preserving researcher oversight and enabling correction of the occasional large-error outliers that would be unacceptable in fully automated workflows. LizardMorph is freely available as an open-source tool and provides a replicable framework for developing ML-assisted annotation tools that can democratize access to high-quality morphometric analysis across diverse biological research communities.

## 1. INTRODUCTION

Morphological measurements form the foundation of much biological research, from ecological field studies to developmental biology, evolutionary research, and conservation monitoring (Adams *et al*. 2004; Mitteroecker & Gunz 2009). Precise quantification of physical traits—body dimensions, skeletal features, organ sizes—enables biologists to identify important population- or species-level patterns, track population health, characterize intra- and interspecific variation, and predict organismal responses to environmental change (Lürig *et al*. 2021). However, obtaining these measurements at the scales required for robust statistical inference is labor-intensive and presents a persistent methodological challenge that constrains the scope and pace of biological research (Fruciano 2016; Lürig *et al*. 2021).

Traditional approaches to morphological measurement fall into two broad categories, each with distinct advantages and limitations. Direct physical measurement of organisms using tools such as calipers, rulers, or other physical measuring devices offers speed and simplicity but introduces substantial measurement error through inconsistent manual placements, observer fatigue, and the physical manipulation of specimens (Fruciano 2016; Yezerinac *et al*. 1992). Digital imaging approaches—analyzing photographs, x-ray radiographs, or computed tomography scans—provide permanent records that can be re-examined and yield highly accurate measurements but require intensive manual labor during processing and data extraction (Devine *et al*. 2020; Lürig *et al*. 2021; Stroud *et al*. 2023, 2024). Researchers must identify and mark dozens of anatomical landmarks on each image using specialized software, a process that typically requires up to several minutes per specimen, even for experienced annotators, depending on landmark complexity (Fruciano 2016). With morphological studies now routinely analyzing hundreds to thousands of specimens, manual landmarking creates a substantial bottleneck that limits research productivity and delays scientific discovery (Devine *et al*. 2020; Lürig *et al*. 2021; Porto & Voje 2020).

The rise of machine learning offers compelling solutions to this measurement bottleneck. Automated landmark detection algorithms can process images orders of magnitude faster than human annotators while maintaining consistent landmark placement criteria (Devine *et al*. 2020; Porto & Voje 2020). Recent advances in computer vision, particularly pose estimation frameworks originally developed for human subjects, have been successfully adapted for biological applications (Mathis *et al*. 2018; Pereira *et al*. 2022). For example, ML-Morph, a landmark detection framework based on ensemble regression trees, has demonstrated remarkably accurate automated landmarking of digital images across diverse taxa (Porto & Voje 2020). As such, machine-learning pipelines like ML-Morph represent substantial technical achievements that present the opportunity to dramatically accelerate biological research (Porto & Voje 2020). However, most automated approaches still require substantial computational expertise for implementation and/or lack the user-friendly interfaces that would make them accessible to biologists without programming backgrounds (Lürig *et al*. 2021). The gap between technical capability and practical usability prevents many research groups from benefiting from automation advances (Lürig *et al*. 2021).

Here, we present LizardMorph, a generalizable machine learning pipeline for automated morphometric landmarking that addresses both technical performance and practical accessibility. Our system comprises two integrated components: a fine-tuned ML-Morph landmark detection model (Porto & Voje 2020), and a web-based interface through which users interact with model predictions. We demonstrate the pipelinès capabilities using lizard radiographs (digital x-ray images) as a case study, where anatomical landmarks can be precisely identified on consistently positioned lizard skeletons (Stroud *et al*. 2023, 2024, 2025; Stroud & Ratcliff 2025). However, LizardMorph‘s architecture is intentionally designed for broader applicability—the underlying framework can be adapted to any biological system where specimens are consistently positioned and imaged.

Critically, LizardMorph implements a human-ML collaboration model rather than pursuing full automation (Kaluarachchi *et al*. 2021). Our web interface displays automated predictions as interactive, editable points overlaid on images, enabling researchers to verify accuracy and correct outlier predictions through intuitive point-and-click interactions common in image annotation workflows. This design philosophy recognizes that biological expertise remains essential for handling edge cases, ambiguous anatomical features, and context-dependent interpretation that automated systems cannot reliably address (Amershi *et al*. 2019; Vella *et al*. 2020). By combining machine efficiency with human oversight, LizardMorph enables researchers to work substantially faster while maintaining the scientific rigor required for publishable research.

Our study has four primary objectives. First, we develop and validate a machine learning pipeline for automated landmark detection on biological images, using lizard radiographs with 34 anatomical landmarks as proof-of-concept. Second, we embed this pipeline within an accessible web interface designed through human-centered design principles, enabling biologists to leverage automation without requiring computational expertise. Third, we evaluate system performance against traditional manual methods across users with varying expertise levels, quantifying both time efficiency and measurement accuracy. Fourth, we assess LizardMorph‘s generalizability to other organisms, imaging modalities, and landmark configurations. By addressing both technical performance and practical usability, LizardMorph provides a replicable framework for developing ML-assisted annotation tools that can democratize access to high-quality morphometric analysis across biological research communities.

In the sections that follow, we explain the computational background and technical details of fine-tuning ML-Morph to optimize it for our needs, describe the LizardMorph web interface and its generalizability as a ‘plug-and-play‘ interface for other ML models, dive into the interfacès performance and evaluation, and conclude by discussing the implications and future adaptations and extensions of our work.

## 2. COMPUTATIONAL BACKGROUND

### 2.1 Case Study: *Anolis* Lizard Digital X-ray Radiographs

Digital X-ray radiographs of *Anolis* lizards (anoles) are collected as part of a long-term field study examining lizard ecology and evolution (Stroud *et al*. 2023, 2024; Stroud & Ratcliff 2025). Each anole in this study is anesthetized, and a dorsal-view radiograph is collected in a standardized position (Figure 1). On each radiograph, 32 anatomical landmarks capture key morphological features of interest to biologists, including cranial dimensions, limb lengths, pelvic girdle structure, and overall body size (Figure 1). Each radiograph includes a 10 mm scale bar for size reference; a landmark is placed at either end, producing a total set of 34 landmarks for each image.

**Figure 1.**
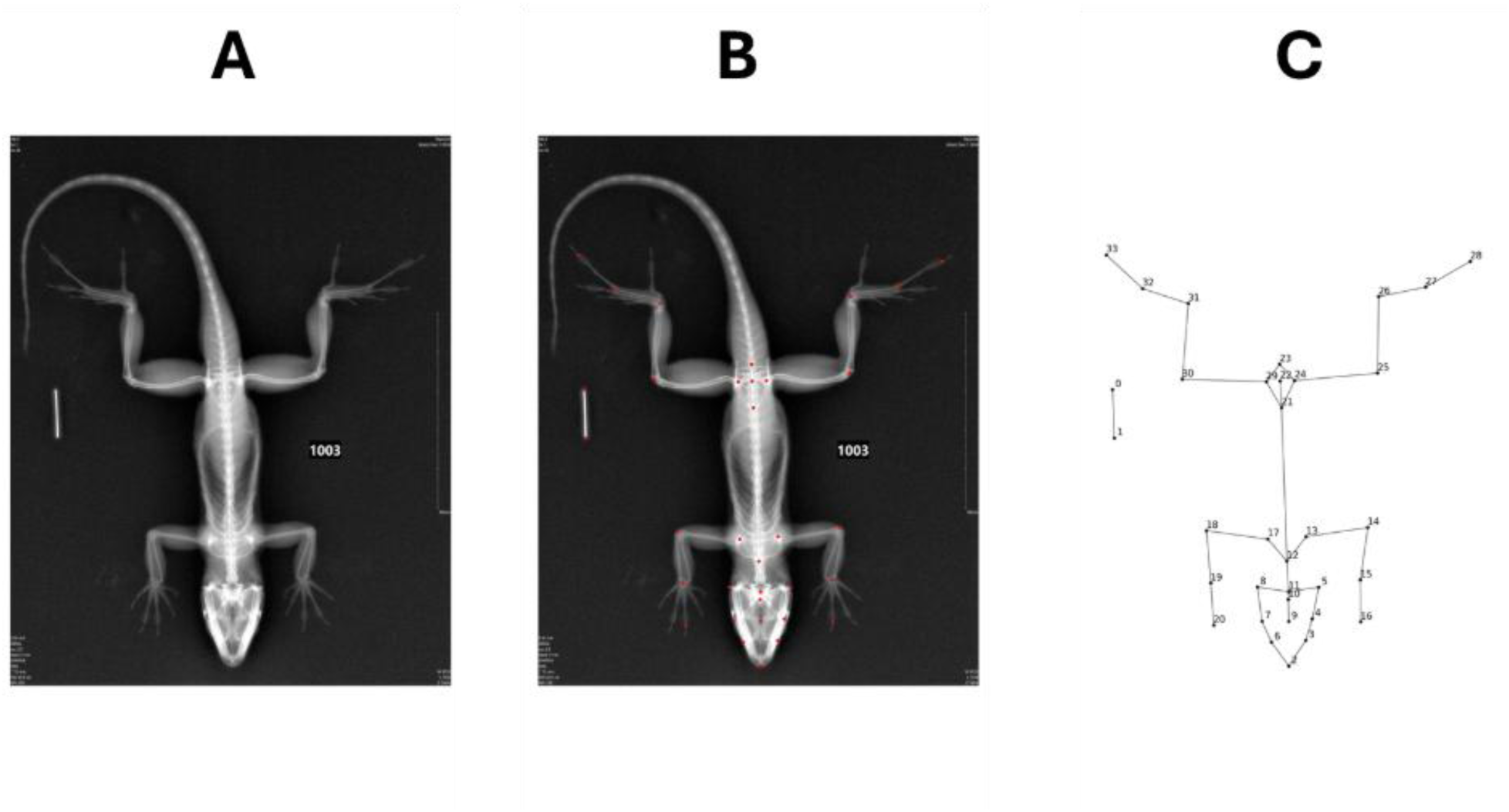
Standardized dorsal radiographs of *Anolis* lizards and the anatomical landmark configuration used for morphometric analysis. (A) An unprocessed dorsal-view digital X-ray radiograph of an *Anolis* lizard (*A. cristatellus*), illustrating the standardized positioning protocol used across all specimens in this study. A 10 mm scale bar (white rectangle, left) is included in each image for size calibration. (B) The same radiograph with all 34 anatomical landmarks overlaid as red points. Landmarks capture key morphological features including cranial dimensions, forelimb and hindlimb lengths, pectoral and pelvic girdle structure, digit tips, and the endpoints of the scale bar. (C) A landmark connectivity diagram showing the skeletal topology defined by the 34-landmark configuration, with landmarks numbered 0–33. This landmark scheme was used for both manual annotation and automated prediction by the LizardMorph ML-Morph model.

These morphological measurements provide critical data for understanding adaptive responses to environmental change, competitive interactions among species, and evolutionary processes shaping lizard communities (Stroud *et al*. 2023, 2024; Stroud & Ratcliff 2025). However, manually placing 34 landmarks on thousands of radiographs—as required in this field study—represents a substantial time investment and introduces potential observer variability that could affect downstream analyses. Automated landmarking approaches offer the potential to dramatically accelerate data collection while maintaining measurement precision, but only if implemented in ways that preserve researcher control over data quality.

### 2.2 Overall Method

LizardMorph is built upon ML-Morph, a pose estimation framework that employs ensemble regression trees to predict anatomical landmark locations on digital images (Porto & Voje 2020). Our implementation follows a two-stage process: first, images undergo preprocessing to enhance anatomical feature visibility; second, a trained shape predictor model identifies and places landmarks based on learned patterns from manually annotated training images. All model training was conducted on a Dell XPS 15 9570 with an Intel Core i7-8750H CPU with 32 GB of RAM using Python 3.11.9 with ‘dlib‘ (Kazemi & Sullivan 2014), ‘opencv-python‘, ‘pandas‘, and ‘numpy‘ (Harris *et al*. 2020). The trained model and web interface are publicly accessible at https://haag-1.cc.gatech.edu/. Source code is available at https://github.com/Human-Augment-Analytics/LizardMorph.

### 2.3 Image Preprocessing and Enhancement

Digital radiographs often exhibit variable contrast that can obscure fine anatomical details critical for accurate landmark placement. To address this challenge, we implemented Contrast Limited Adaptive Histogram Equalization (CLAHE) using OpenCV (Zuiderveld 1994). CLAHE was selected over standard histogram equalization because it enhances local contrast within defined image regions while preventing noise amplification in uniform soft tissue areas (Zuiderveld 1994). For our radiographic images, this tile-based approach ensures that both large skeletal structures—such as the limbs, skull, and pelvis—and fine anatomical details—such as digit bones—receive appropriate contrast enhancement without over-saturating the entire image. This preprocessing step significantly improved the visibility of skeletal boundaries, which is critical for accurate landmark detection (Figure 4).

### 2.4 Training Data Preparation

We prepared training data by manually annotating all 34 anatomical landmarks on 838 lizard radiographs. All manual annotations were performed by a single experienced observer (J.J.S.) to maintain consistency in landmark placement criteria. Following annotation and CLAHE preprocessing, the dataset was randomly partitioned into training (80%, n = 670 images) and testing (20%, n = 167 images) subsets to enable independent validation of model performance.

### 2.5 Model Training

The standard ML-Morph pipeline includes an object detection step to localize subjects within images before landmark prediction (Porto & Voje 2020). However, our radiographic protocol ensures consistent specimen positioning and orientation across all images, with lizards centered in the frame in a standardized dorsal view (Figure 1). This consistency allowed us to eliminate the object detection stage, focusing computational resources entirely on landmark precision rather than subject localization. By removing this step, we simplified the pipeline while improving processing speed without sacrificing accuracy.

We trained the shape predictor using the following parameters: tree depth of 5, cascade depth of 35, test split of 20%, regularization parameter of 0.08, oversampling amount of 300, 500 regression trees, and feature pool size of 1200. These parameters were selected through preliminary optimization to balance prediction accuracy with computational efficiency. The cascade depth determines how many stages of refinement the model applies to landmark predictions, with deeper cascades generally improving accuracy at the cost of longer training and inference times (Porto & Voje 2020). The regularization parameter controls model complexity to prevent overfitting to training data, while the oversampling amount determines how many random perturbations of training images the model encounters, improving robustness to natural variation in image quality and specimen positioning (Porto & Voje 2020).

#### 2.5.1 Parameter Optimization

Hyperparameters for the landmark prediction model were tuned using the ‘find_min_global‘ routine in ‘dlib‘ (Kazemi & Sullivan 2014). This is a derivative-free global optimizer for bounded black-box functions. The method is based on the LIPO family of global optimization algorithms for Lipschitz functions and, in ‘dlib‘, is implemented as a hybrid procedure that alternates between global exploration using an upper-bound model of the objective function and local refinement using a trust-region search (Malherbe & Vayatis 2017). This strategy is well-suited to hyperparameter tuning because it does not require gradients or manually chosen starting values and is designed to use relatively few objective evaluations (Koch *et al*. 2018).

For each candidate hyperparameter configuration, we trained an ML-morph landmark predictor on the training set and evaluated its performance on a held-out validation set using normalized landmark error. The search was performed over predefined bounds for tree depth, cascade depth, test split size, regularization parameter, oversampling amount, number of regression trees, and feature pool size. Optimization proceeded for 100 objective function evaluations, and the final hyperparameter set was chosen as the configuration that minimized validation error. The final model was then retrained using these selected parameters on training data and evaluated on the independent test set.

### 2.6 Model Performance Evaluation

We evaluated model performance on the held-out test set (n = 167 images) by calculating the Euclidean distance between each predicted landmark and the corresponding expert-placed landmark from the original manual annotations. To convert pixel-based distances into biologically meaningful units, we used the 10 mm scale bar present in each radiograph to transform measurements into millimeters. Following established morphometric accuracy standards (Porto & Voje 2020), we classified predictions as acceptable if they fell within 1 mm of the expert landmark location.

Across all 34 landmarks and all test images, LizardMorph achieved 82% accuracy overall, with predictions falling within the 1 mm tolerance threshold (Figure 2). Prediction accuracy varied systematically across landmark types. Landmarks located on large, well-defined skeletal structures with consistent positioning—such as major limb joints, pelvic landmarks, and cranial features—exhibited the highest accuracy (100%). In contrast, landmarks associated with digit tips and the scale bar showed lower accuracy (86%) and occasional large-error outliers exceeding 3 mm. We attribute this reduced performance to spatial inconsistency: digit positions vary substantially depending on how specimens are positioned during radiography, and scale bars appear in different image locations across photographs. These spatially inconsistent features provide a less reliable training signal compared to landmarks that appear in stereotyped locations across specimens.

**Figure 2.**
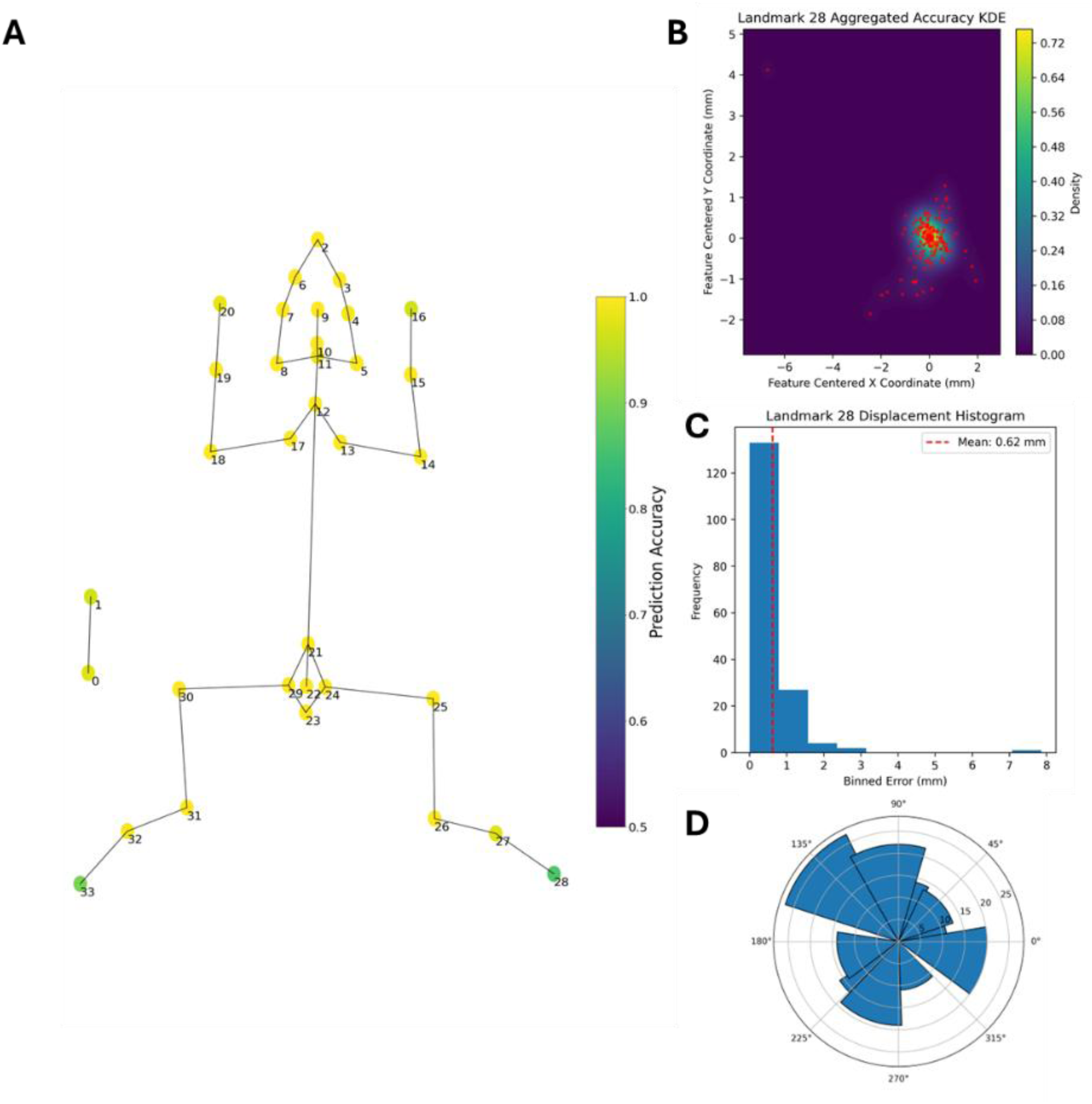
LizardMorph model prediction accuracy across all 34 landmarks, with detailed diagnostics for the lowest-performing landmark. (A) Landmark connectivity diagram showing prediction accuracy across all 34 landmarks aggregated over the held-out test set (n = 167 images). Each landmark node is colored according to the proportion of test images in which its predicted location fell within the 1 mm tolerance threshold. The majority of landmarks achieved 100% accuracy; landmarks 28 and 33 (green) showed the lowest accuracy at ca. 86%. (B) Kernel density estimation (KDE) of predicted landmark locations for Landmark 28, centered on the expert-placed ground truth position. Axes represent displacement in millimeters from the true landmark location. The dense cluster near the origin confirms that most predictions are highly accurate, while scattered red points indicate occasional large-error outliers. (C) Displacement histogram for Landmark 28, showing the distribution of Euclidean errors (mm) across all test images. The mean displacement was 0.62 mm (red dashed line), reflecting overall good predictive performance even for the worst-performing landmark prediction. (D) Polar histogram showing the directional distribution of prediction errors for Landmark 28. The roughly uniform distribution across all angles indicates that errors are not systematically biased in any anatomical direction, suggesting that outlier errors reflect stochastic positional variability rather than consistent model bias.

Following successful validation of ML-Morph model‘s performance on lizard radiographs, we next designed a web interface to make this automated landmarking accessible to biologists without computational expertise.

## 3. LizardMorph: WEB INTERFACE DESIGN AND WORKFLOW

### 3.1 Design Philosophy and User-Centered Development

Following human-centered design principles, we embedded the trained model within a user interface designed to facilitate its use without requiring machine learning or computational expertise (Amershi *et al*. 2019). Our design process was a close collaboration between the development team (comprised of computer scientists) and the end users (lizard biologists), who informed the necessary features of the tool through iterative prototype testing and feedback on workflow efficiency and interface design. This collaborative approach guided the evolution from a single-image interface to a comprehensive batch processing system that maintains familiar interaction patterns while integrating automation capabilities.

We designed the interface around human-ML collaboration as the foundational principle for effective scientific annotation tools (Wang *et al*. 2019). Our interface design implements three human-AI interaction guidelines (Amershi *et al*. 2019): (1) Decision Transparency, making ML predictions transparent and interpretable through visualization; (2) Interactive Correction, enabling users to directly edit and refine ML predictions (Fails & Olsen 2003); and (3) Simple Global Control, providing straightforward system-wide settings that welcome participation at all skill levels (Amershi *et al*. 2019).

Our goal was to create a tool that enhances rather than replaces scientific expertise. While automated analysis may not be perfect, it can democratize access to high-quality morphometric analysis while maintaining the scientific rigor required for research (Porto & Voje 2020). Furthermore, such systems address a persistent challenge in research laboratories: the constant need to retrain new team members in complex analysis techniques. As one of our collaborators noted regarding manual landmarking workflows: the ability to onboard new researchers efficiently while maintaining analytical standards represents a significant advancement for the sustainability and scalability of ecological research programs.

### 3.2 System Architecture and Workflow

LizardMorph provides a streamlined workflow for anatomical landmarking through a web-based application accessible via standard browsers without requiring local software installation (Figure 3). The system consists of a frontend user interface and a backend server hosting the trained detection and classification models. Users begin by uploading a batch of up to 100 JPEG images through the web interface. The system automatically processes these images using the pre-trained LizardMorph model to predict landmark locations. These predictions are displayed as interactive, editable points overlaid directly on each image, allowing users to manually review and adjust their positions for accuracy. Once satisfied, users can download a single ZIP archive containing all predictions and outputs (see Section 3.4 for full details). The system is deployed on Georgia Tech‘s College of Computing server and accessible at [https://haag-1.cc.gatech.edu/].

**Figure 3.**
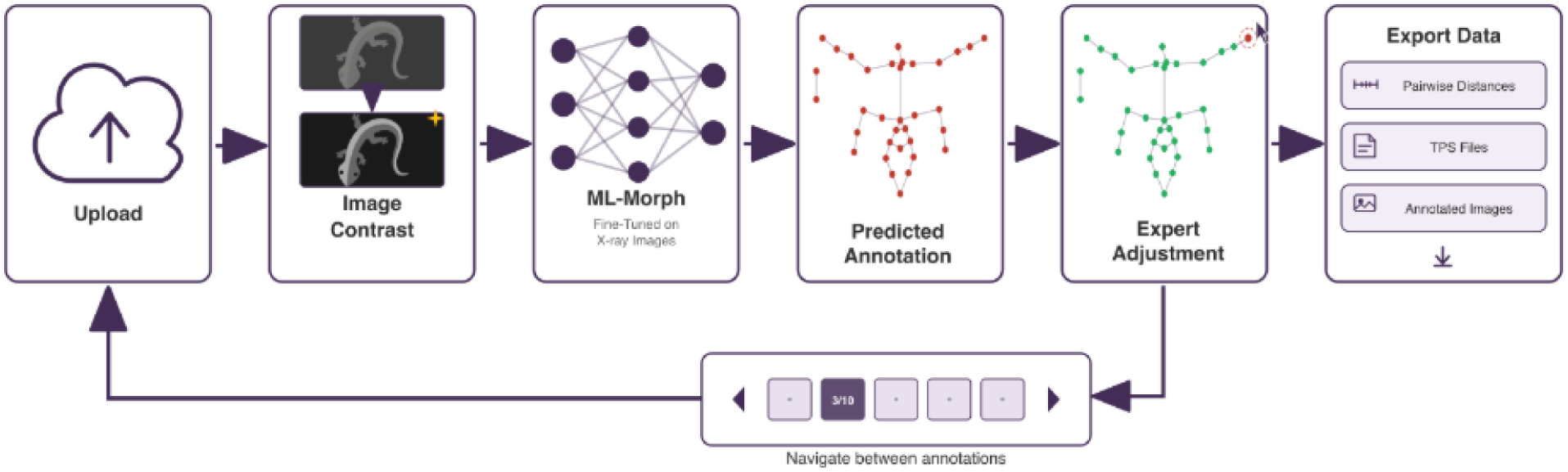
The LizardMorph processing pipeline, illustrating the six-stage workflow from image upload to data export. (1) Upload: Users submit batches of up to 100 JPEG radiographs through the web interface. (2) Image Contrast: Each image undergoes automated preprocessing, including Contrast Limited Adaptive Histogram Equalization (CLAHE), to enhance skeletal boundary visibility prior to landmark detection. (3) ML-Morph: The fine-tuned ML-Morph shape predictor model, trained on lizard radiographs, automatically places 34 anatomical landmarks on each preprocessed image. (4) Predicted Annotation: Automated landmark predictions (red points) are displayed on the radiograph as an interactive overlay, ready for expert review. (5) Expert Adjustment: Users inspect and manually correct any inaccurate predictions by clicking and dragging landmarks to their true anatomical positions (corrected landmarks shown in green). A batch navigation panel beneath the workflow allows users to move sequentially between all uploaded images in the session. (6) Export Data: Once all images are reviewed, users download a ZIP archive containing three output types: pairwise landmark distance matrices (CSV), landmark coordinates in TPS file format compatible with standard geometric morphometrics software, and annotated JPEG images showing final landmark placements for visual verification.

### 3.3 Interface Components

The interface is organized into three primary components: the Control Panel, the History Panel, and the SVG Viewer (Figure 4). These components work together to provide an efficient workflow for processing multiple images while maintaining user control over prediction quality.

#### 3.3.1 Control Panel

The Control Panel serves as the central hub for session management and data operations (Figure 4). Users initiate or continue a session by clicking the “Upload X-Ray Images” button, which triggers the file selection dialog. The panel accepts JPEG format images with file sizes up to 10 MB per image. Once images are selected, they are transmitted to the server for automated processing. The “Export All Data” button initiates a ZIP file download containing all model predictions in TPS file format, a CSV containing all pre-calculated pairwise landmark distances to eliminate manual post-processing, and visualizations of each image with predicted landmarks overlaid (see Section 3.4). The “Clear History” button clears the current session and deletes all images and predictions from browser memory, allowing users to begin new analysis sessions.

#### 3.3.2 History Panel

The History Panel displays uploaded images with real-time processing status indicators (Figure 4). For each file being processed, a dedicated progress bar provides visual feedback tracking the status of both file upload and the subsequent automated landmarking pipeline. The progress bar displays specific completion stages: 25% indicates the start of file upload, 50% represents successful file transfer to the server, 75% shows completion of image preprocessing and landmark detection, and 100% signifies full processing completion with all data ready for review. Upon completion, each image is marked as ready along with a timestamp, enabling users to monitor the status of large batches and select any completed image for immediate review. This real-time feedback mechanism enables users to understand system status and manage their workflow efficiently.

**Figure 4.**
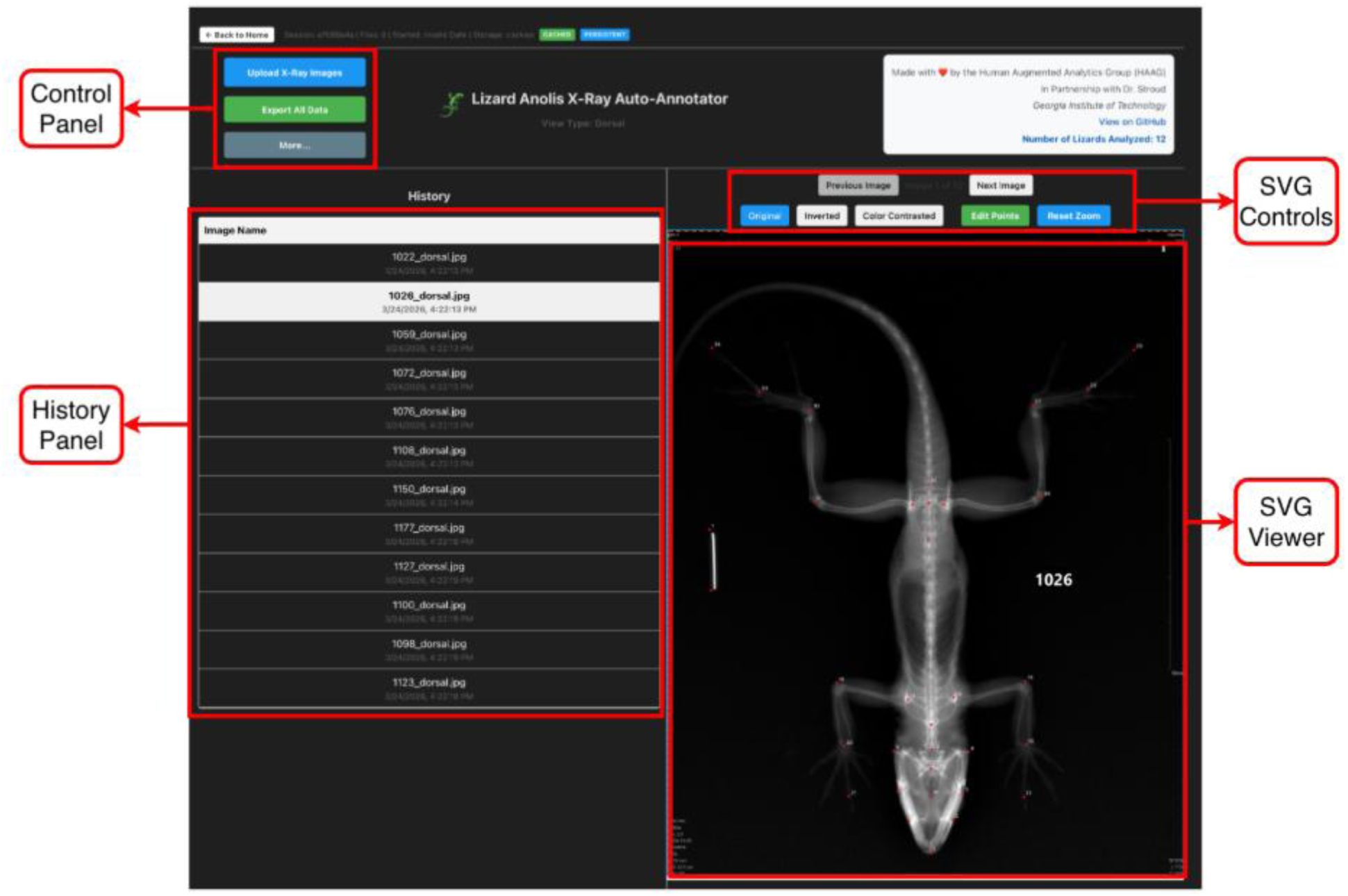
The LizardMorph web interface, showing the three primary components used for batch landmark annotation. (1) Control Panel (top left): Contains three primary action buttons — “Upload X-Ray Images” to initiate or add to a batch session, “Export All Data” to download all predictions and outputs as a ZIP archive, and “More…” for additional session settings. A session status bar displays file count and storage information. (2) History Panel (left): Displays the full list of uploaded images by filename and timestamp, allowing users to navigate between specimens by clicking any entry. The currently active image (here, 1026_dorsal.jpg) is highlighted, and the panel tracks all images processed within the session. (3) SVG Controls and Viewer (right): The SVG Controls toolbar provides buttons to toggle between three contrast modes — Original, Inverted, and Color Contrasted — as well as “Edit Points” to activate manual landmark correction, “Reset Zoom” to restore default magnification, and “Previous Image”/”Next Image” for sequential navigation. The SVG Viewer displays the active radiograph with automated landmark predictions overlaid as interactive red points, shown here on specimen 1026 in the default Original contrast mode. Together, these components enable efficient batch review and correction of automated landmark predictions without requiring any local software installation.

#### 3.3.3 SVG Viewer

The SVG Viewer provides the core interaction space for landmark inspection and correction, displaying the selected radiograph with predicted landmarks overlaid as interactive points (Figure 4). The viewer implements several features to aid precise landmark placement and verification:

##### Multiple Viewing Modes

For each input image, LizardMorph creates three enhanced versions to provide researchers with multiple perspectives for landmark validation (Figure 5). This multi-modal approach addresses the inherent variability in X-ray image quality and lighting conditions. Users can toggle between three distinct image representations: (1) Original—the unmodified X-ray for baseline reference and comparison; (2) Inverted—color complement calculated as

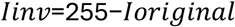

**Figure 5.**
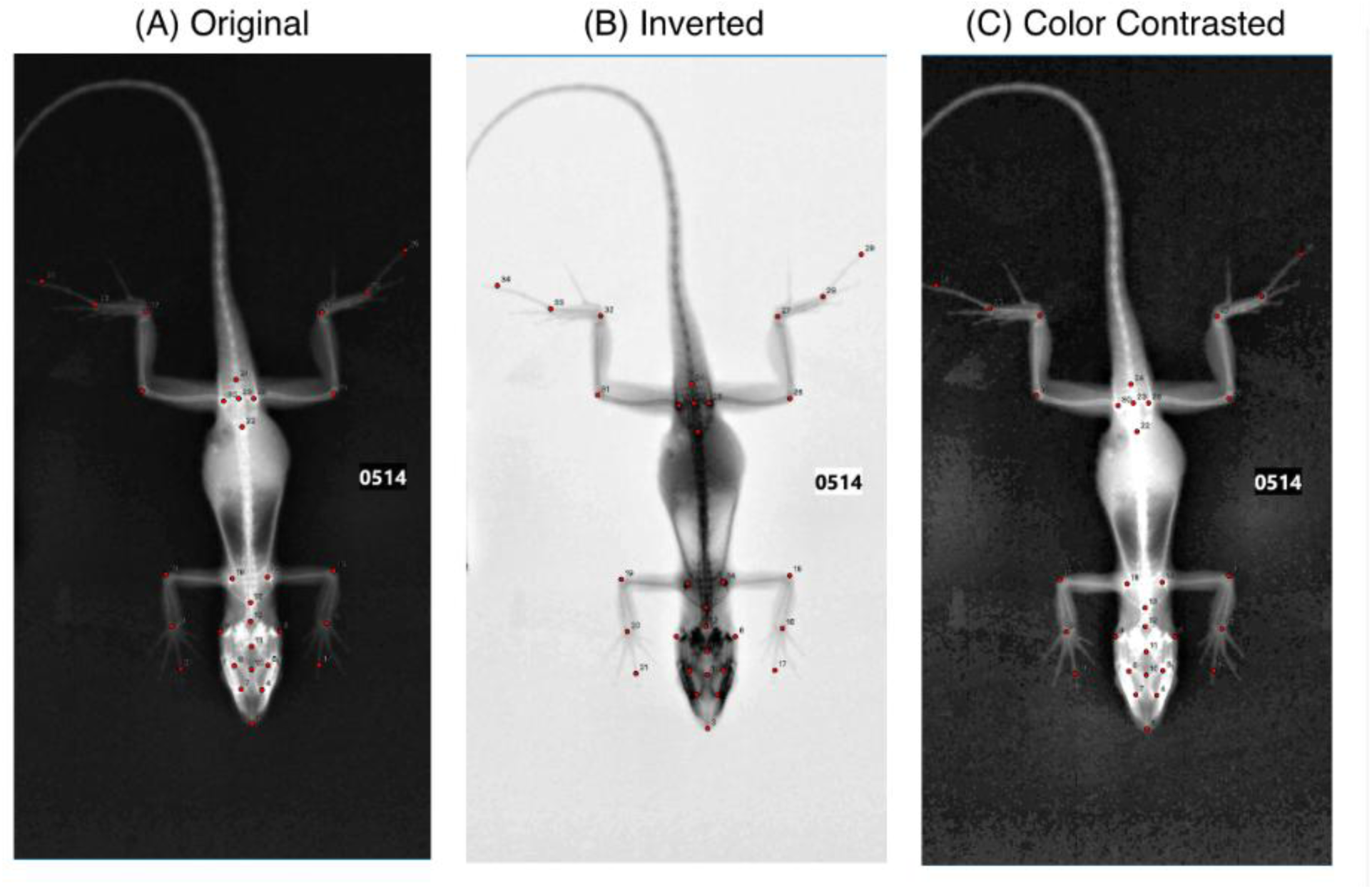
The three image contrast modes available in the LizardMorph SVG Viewer, demonstrated on specimen 0514 with all 34 predicted landmarks overlaid as red points. (A) Original: The unmodified grayscale radiograph, providing a baseline view of specimen anatomy. Bone density appears brightest in regions of dense skeletal tissue such as the pelvic girdle and limb joints. (B) Inverted: The color-complement transformation (I_inv = 255 − I_original), which reverses the grayscale values so that dense structures appear dark against a light background. This mode improves visibility of landmarks positioned over high-density skeletal regions where red points may be difficult to distinguish in the original view. (C) Color Contrasted: The CLAHE-processed image used during model training, which enhances local contrast to improve visibility of fine skeletal structures such as digit bones and cranial features. Users can toggle between all three modes during landmark review to confirm accurate placement across varying anatomical regions and image qualities.

which enhances visibility of certain anatomical features that may be obscured in the original grayscale; and (3) Enhanced—the CLAHE-processed image described in Section 2.3, which provides superior visualization of fine skeletal details. These viewing modes are crucial for ensuring landmark accuracy across different image qualities and lighting conditions.

##### Interactive Landmark Correction

The “Edit Points” mode enables users to manually click and drag any landmark to a corrected position (Figure 6). This interactive correction capability is crucial for ensuring dataset quality, as it allows expert refinement of the initial automated predictions. When edit mode is activated, zoom and pan functionality is temporarily disabled to prevent accidental navigation during precise point placement. Selected points are highlighted in yellow, while unselected points remain red, providing clear visual feedback during editing. Users can save corrected positions or cancel edits to revert to automated predictions.

**Figure 6.**
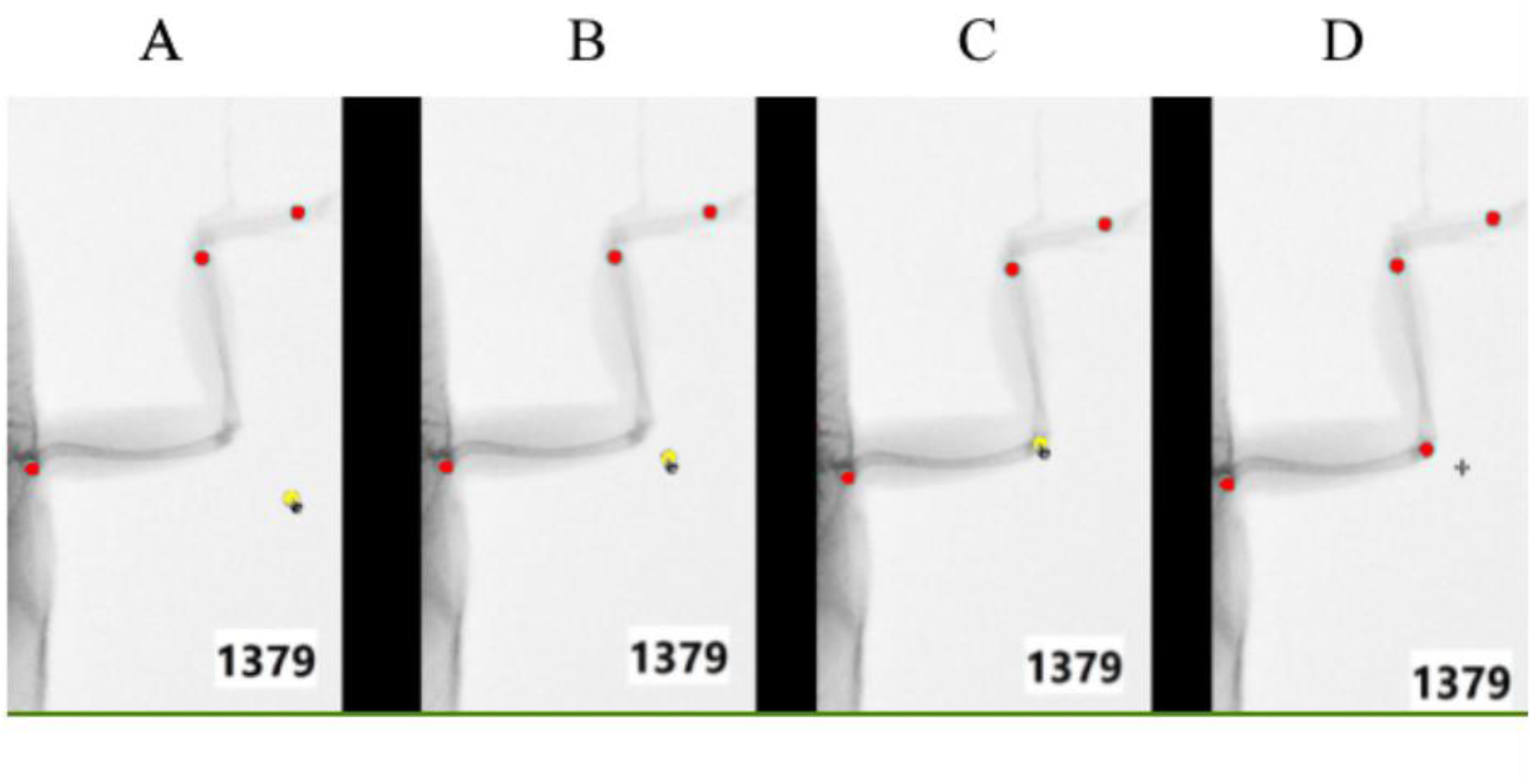
The interactive landmark correction workflow in LizardMorph, demonstrated on a zoomed-in forelimb region of specimen 1379. When a user activates “Edit Points” mode, individual landmarks can be selected and repositioned by clicking and dragging. (A) Initial state: automated predictions displayed as red points, with one landmark (shown in yellow) selected for correction. (B–C) The selected landmark is actively being dragged toward its true anatomical position; the yellow highlight indicates the actively selected point throughout the editing process, while unselected landmarks remain red. (D) The corrected landmark has been placed at its true anatomical position by the user, with the point returning to red upon deselection. This click-and-drag correction workflow enables rapid expert refinement of outlier predictions while preserving accurate automated placements elsewhere in the image.

##### Navigation and Visualization Controls

Standard zoom-in and zoom-out functions are available for close inspection of anatomical details, with zoom levels ranging from 0.5x to 5x magnification. The zoom behavior is implemented using D3.js transformations, ensuring smooth and responsive interactions (Bostock 2011). The “Reset Zoom” button returns the image to its default view (1x magnification) and centers the image within the viewer. Additional visualization controls allow users to adjust landmark point size and transparency to optimize visibility against different image backgrounds (Figure 4).

### 3.4 Data Export and Session Management

Once users are satisfied with landmark placements for all uploaded images, they can export all data through a single action via the “Export All Data” button. The system compiles all session data into a ZIP file containing three types of output: (1) TPS files with landmark coordinates in the standard format used by geometric morphometrics software (e.g., MorphoJ; (Klingenberg 2011) or specialists R packages (e.g., ‘geomorph‘; (Adams & Otárola-Castillo 2013), with separate files containing both original automated predictions and user-corrected coordinates to ensure data integrity and traceability; and (2) annotated JPEG images showing the original radiograph with landmark positions overlaid, providing visual documentation of landmark placement for quality control and publication; and (3) a CSV file containing pre-calculated pairwise landmark distances, eliminating the need for manual post-processing before downstream morphometric analysis. This three-part output format serves both quantitative analysis needs and qualitative verification requirements.

All processing occurs server-side, with results temporarily stored in the user‘s browser session, ensuring that no data persists on servers after the session ends—an important consideration for researchers working with unpublished or sensitive data.

### 3.5 Generalizability

LizardMorph‘s framework extends beyond lizard radiographs: any biological imaging workflow in which specimens are consistently positioned can be supported by training an ML-Morph predictor on a new dataset and deploying it through the interface without modification. To demonstrate this, we trained an ML-Morph model on the Drosophilid wing dataset from Porto & Voje (2020), comprising 280 microscopy images of fruit fly wings with 12 landmarks at wing vein intersections, and integrated it directly into the LizardMorph web application, allowing users to upload wing images and interact with predicted landmarks through the same editing and export workflow used for lizard radiographs.

## 4. PERFORMANCE EVALUATION

### 4.1 Study Design and Objectives

To evaluate whether LizardMorph achieves its goals of improving efficiency while maintaining accuracy, we conducted a controlled user study comparing our system against traditional manual methods. Our evaluation focused on two critical dimensions: efficiency (task completion time) and accuracy (measurement precision), as both are essential for practical adoption in scientific workflows. Additionally, we conducted A/B testing to optimize interface design elements before evaluating overall system performance.

We recruited participants representing two distinct levels of expertise: (1) an experienced annotator—a researcher with extensive background in morphometric analysis and familiarity with TpsDig2 software (Rohlf 2015), having previously annotated thousands of lizard radiographs; and (2) an inexperienced annotator—a student new to landmark annotation with minimal prior experience, representing typical new laboratory members who require training in morphometric methods.

### 4.2 A/B Testing: Interface Optimization

To optimize our interface design before evaluating system performance, we conducted A/B testing comparing two landmark visualization approaches: Transparent Mode, where landmark points are rendered as semi-transparent circles allowing underlying anatomical features to remain visible, and Outline Mode, where landmarks are displayed as outlined circles with hollow centers (Figure 7).

**Figure 7.**
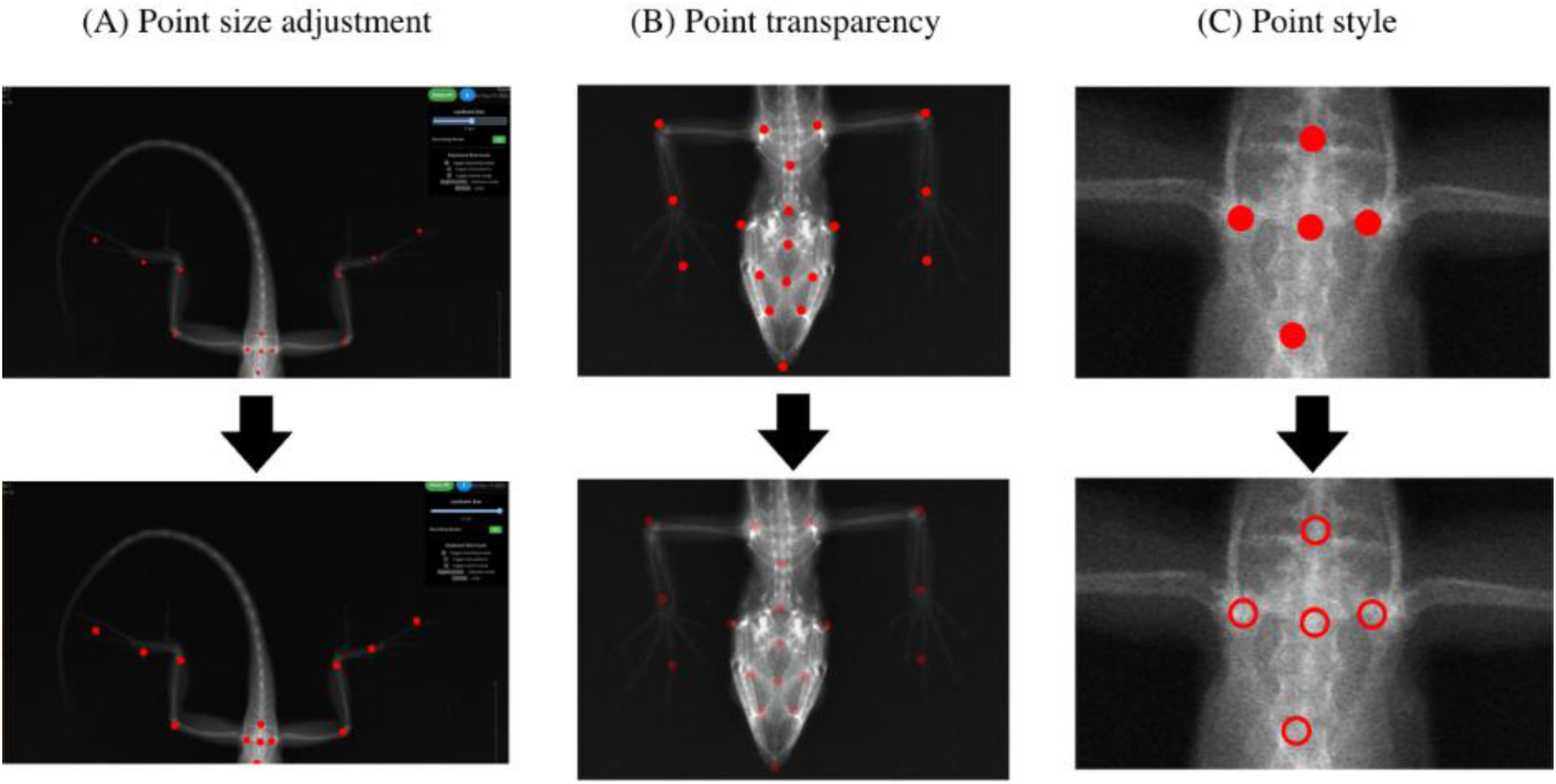
Landmark visualization customization controls in LizardMorph. (A) Point size adjustment: larger landmarks (6.0 px, bottom) provide high visibility across the full specimen, while smaller landmarks (3.0 px, top) reduce occlusion in densely landmarked regions such as the pelvic girdle and digit joints. (B) Point transparency: solid landmarks (top) are maximize point visibility, while semi-transparent landmarks (bottom) allow underlying skeletal structures to show through — useful when verifying placement on fine or overlapping features. (C) Point style: filled circles (top) offer high contrast against dark backgrounds, while hollow outline circles (bottom) minimize obstruction of underlying anatomy. Together these controls allow users to optimize landmark visibility for different image qualities and anatomical regions.

#### 4.2.1 Methods

Our experienced annotator landmarked identical images using both visualization modes in randomized order, allowing us to control individual differences and image complexity. We randomly selected 5 test images representing a range of specimen sizes and image quality, and measured completion times for landmarking tasks under each visualization condition.

#### 4.2.2 Results

The A/B testing results demonstrate that Transparent Mode consistently outperformed Outline Mode across test images, achieving an average time improvement of 18.9% (Table 1). The most substantial improvement on a single image reduced completion time by 43.6% (from 1:18 to 0:44). These findings suggest that transparent landmark visualization provides better visual clarity for rapid landmark identification and placement. The transparent approach allows users to simultaneously view both the underlying anatomical structures and the landmark positions without visual obstruction, leading to more efficient annotation workflows. Based on these results, we selected Transparent Mode as the default visualization for all subsequent performance evaluations and system deployment.

**Table 1.**
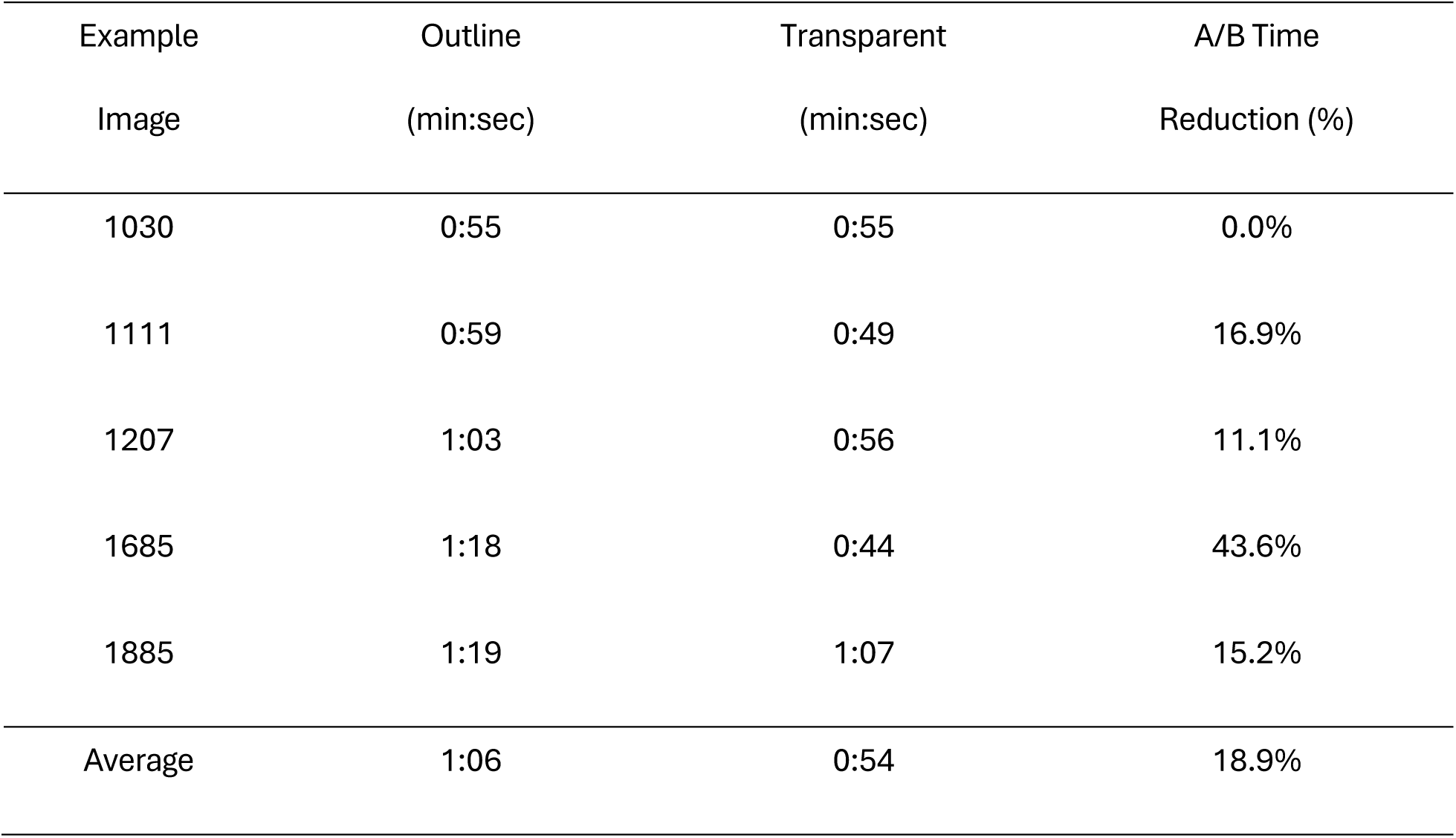
Interface optimization and performance comparison for 5 random images. Left: A/B testing results comparing Outline Mode vs Transparent Mode visualization for experienced users across 5 test images. Right: LizardMorph performance using optimized Transparent Mode interface by an ‘experienced landmarker’ compared to traditional manual annotation methods, showing substantial time reductions across all specimens.

### 4.3 Performance Comparison: LizardMorph vs. Manual Methods

#### 4.3.1 Experimental Design

The study used a within-subjects design, where each participant annotated the same 10 lizard X-ray images using traditional software (TpsDig2) and then again using LizardMorph with the optimized Transparent Mode interface. For each image, participants were asked to place all 34 landmarks according to standardized anatomical definitions. We measured completion time from the moment an image was opened until all landmarks were placed and saved. The 10 specimens were drawn at random from the held-out test set, and both annotators processed the same image set to enable direct comparison across expertise levels.

#### 4.3.2 Efficiency Results

Our evaluation showed that LizardMorph produced a clear efficiency benefit for the experienced annotator, while the inexperienced annotator did not show a timing benefit on first exposure (Table 2). During the experiment, the inexperienced annotator was using both software tools for the first time and moved each landmark to verify the correct placement. We interpret this behavior as part of the normal learning process associated with novice performance and increased cognitive effort during skill acquisition (Persky & Robinson 2017). The experienced annotator completed the 10-image set in 6m35s (395 s) with LizardMorph compared with 10m32s (632 s) in TpsDig2, a savings of 3m57s (237 s; 37.5%). LizardMorph was faster on every image, and the paired reduction was statistically significant (Wilcoxon signed rank: W = 0, Z = -2.75, p < 0.005).

**Table 2.** Per-image annotation times for all 10 test specimens, for both expertise levels and both tools. Reduction (%) is (TpsDig2 - LizardMorph) / TpsDig2, so positive values indicate LizardMorph was faster.

| Lizard ID | Exp.<br>TpsDig2 | Exp.<br>LizardMorph | Exp.<br>Reduction | Inexp.<br>TpsDig2 | Inexp.<br>LizardMorph | Inexp.<br>Reduction |
| --- | --- | --- | --- | --- | --- | --- |
| 1029 | 0:57 | 0:43 | 24.6% | 4:33 | 4:55 | -8.1% |
| 1048 | 1:01 | 0:38 | 37.7% | 4:02 | 3:46 | 6.6% |
| 1058 | 1:09 | 0:52 | 24.6% | 3:16 | 2:41 | 17.9% |
| 1063 | 1:04 | 0:55 | 14.1% | 2:32 | 5:18 | -109.2% |
| 1074 | 1:01 | 0:24 | 60.7% | 2:13 | 1:11 | 46.6% |
| 1096 | 1:03 | 0:31 | 50.8% | 2:53 | 2:14 | 22.5% |
| 1111 | 1:15 | 0:58 | 22.7% | 2:34 | 4:48 | -87.0% |
| 1132 | 1:04 | 0:29 | 54.7% | 2:11 | 2:57 | -35.1% |
| 1146 | 0:59 | 0:22 | 62.7% | 2:36 | 2:56 | -12.8% |
| 1189 | 0:59 | 0:43 | 27.1% | 2:11 | 3:21 | -53.4% |
| Mean | 1:03 | 0:40 | 37.5% | 2:54 | 3:25 | -17.6% |
| Total | 10:32 | 6:35 | 37.5% | 29:01 | 34:07 | -17.6% |

#### 4.3.3 Accuracy Results

We assessed measurement accuracy by comparing all experimental annotations against the experienced TpsDig2 set, which served as our manual reference. We utilized two primary metrics: pairwise landmark-distance error, which summarizes differences in exported morphometric measurements (Adams *et al*. 2004; Mitteroecker & Gunz 2009), and landmark-coordinate error, which measures point placement precision. All errors were converted to millimeters using the 10 mm scale bar.

These results show that LizardMorph preserved accuracy while supporting faster expert annotation. Against the experienced TpsDig2 reference, experienced LizardMorph annotations had a pairwise-distance MAE of 0.388 mm and a landmark-coordinate MAE of 0.212 mm; inexperienced LizardMorph annotations had MAEs of 0.482 mm and 0.253 mm, respectively; and inexperienced TpsDig2 annotations had MAEs of 0.769 mm and 0.336 mm, respectively. The novice comparison was especially informative: despite spending a similar total amount of time in the two workflows, the inexperienced annotator‘s TpsDig2 measurements were substantially less accurate than their LizardMorph measurements, with pairwise-distance error increasing from 0.482 mm in LizardMorph to 0.769 mm in TpsDig2.

Thus, LizardMorph did not simply trade precision for speed. For experienced users, it reduced processing time while maintaining measurement agreement; for inexperienced users, it appeared to scaffold more consistent measurement placement even before producing a timing advantage.

## 5. DISCUSSION

The manual landmarking workflows on which most morphometric studies depend remain time- and labor-intensive bottlenecks that limit both the pace and scale of ecological and evolutionary research (Fruciano 2016; Lürig *et al*. 2021). Here, we presented LizardMorph, an integrated machine learning pipeline and web-based interface for semi-automated anatomical landmark detection on biological images. Using dorsal radiographs of *Anolis* lizards annotated with 32 anatomical landmarks (plus two scale-bar landmarks) as a proof-of-concept, we addressed four objectives: (1) developing and validating an ML-Morph-based landmark detection model; (2) embedding this model within an accessible, human-centered web interface; (3) evaluating system performance against traditional manual methods across users with varying expertise; and (4) demonstrating generalizability to alternative organisms and imaging modalities. Our results show that LizardMorph substantially reduces annotation time—particularly for experienced users—while maintaining measurement accuracy equivalent to expert manual methods, and that the underlying framework generalizes beyond its original application.

LizardMorph differs from prior automated landmarking tools in its deliberate integration of two components that are typically developed in isolation: a domain-specific ML model and an intuitive graphical interface that supports human oversight. While ML-Morph (Porto & Voje 2020) demonstrated that ensemble regression trees can achieve high landmark accuracy across diverse taxa, its command-line implementation requires programming proficiency that excludes many potential users in ecology and evolution. Conversely, established annotation platforms such as TpsDig2 (Rohlf 2015) provide accessible interfaces but rely entirely on manual placement. LizardMorph bridges this gap by coupling automated prediction with interactive, point-and-click correction within a browser-based environment, implementing a human-in-the-loop collaboration model (Kaluarachchi *et al*. 2021). This design philosophy recognizes that full automation is neither achievable nor desirable for high-precision morphometric research: biological expertise remains essential for interpreting ambiguous anatomical features, handling edge cases, and ensuring data quality (Vella *et al*. 2020).

### 5.1 ML Model Performance and the Value of Image Preprocessing

The LizardMorph ML-Morph model achieved high overall predictive accuracy across all 34 landmarks, with most predictions falling within the 1 mm tolerance threshold established in prior morphometric work (Figure 2; (Porto & Voje 2020)). Performance varied systematically by landmark type: landmarks on large, well-defined skeletal structures — such as major limb joints, pelvic landmarks, and cranial features — achieved 100% accuracy, whereas digit tips and scale bar endpoints exhibited lower accuracy (Figure 2). This is consistent with the principle that pose estimation models perform best when target features occupy predictable spatial positions (Mathis *et al*. 2018; Pereira *et al*. 2022): digit tip positions vary with specimen handling and scale bar locations vary with imaging setup, both providing a weaker and more variable training signal than anatomically fixed landmarks.

The CLAHE preprocessing step contributed meaningfully to model performance by enhancing local contrast across defined image regions without amplifying noise in uniform soft-tissue areas (Zuiderveld, 1994), improving visibility of skeletal boundaries critical for accurate landmark detection. Beyond lizard radiographs, CLAHE is broadly applicable to biological imaging contexts where contrast variability poses challenges, including histological sections, micro-CT reconstructions, and field photography under variable lighting (Mohd-Isa et al., 2021). Similarly, eliminating the object detection stage standard in the ML-Morph pipeline (Porto & Voje, 2020) — justified by our standardized radiographic protocol — improved processing speed and likely contributed to high accuracy on spatially consistent landmarks. Both decisions illustrate how targeted modifications to an existing pipeline can yield meaningful performance gains, though they also mean LizardMorph is currently optimized for imaging workflows with consistent specimen positioning — a constraint to consider when adapting the framework to systems with greater positional variability.

### 5.2 Accessibility Through Human-Centered Design

A central design goal of LizardMorph was to make automated landmarking accessible to researchers without computational expertise, addressing a well-documented gap between ML capability and practical adoption in biological research. Through iterative collaboration between computer scientists and biologists, the resulting interface requires no local installation, command-line interaction, or familiarity with machine learning: users upload images, review overlaid predictions, correct misplaced landmarks by dragging, and export results — all within a standard web browser.

The human-in-the-loop design serves as a critical scientific function beyond usability. Although automated predictions provide excellent starting points for most landmarks (median error ∼0.19 mm), uncorrected predictions include occasional large errors exceeding 3 mm that would introduce unacceptable noise into downstream analyses. Interactive correction transforms this limitation into a practical strength: researchers verify predictions, correct the small proportion of outliers, and produce datasets of equivalent quality to fully manual annotation in substantially less time — demonstrating that biological expertise and automated efficiency are complementary rather than competing resources (Wang *et al*. 2019). Additionally, by providing automated predictions as scaffolding, LizardMorph reduces the learning curve for new personnel, enabling students and technicians to contribute productively to accurate data collection more quickly — a benefit consistently identified as valuable by the biologists (J.J.S., J.T.S.) during the design process.

### 5.3 Efficiency and Accuracy Across Expertise Levels

Our performance evaluation demonstrated that LizardMorph yields substantial efficiency gains while maintaining accuracy equivalent to traditional manual methods. Experienced users achieved a 37.5% reduction in annotation time compared to TpsDig2, which was statistically significant (Wilcoxon Signed-Rank test, W = 0, p < 0.005). Inexperienced users did not show a significant time reduction on first exposure, but their measurements were more accurate in LizardMorph than in TpsDig2 over a similar session length (pairwise-distance error: 0.482 mm vs. 0.769 mm; landmark-coordinate error: 0.253 mm vs. 0.336 mm).

The benefit to experienced users is particularly noteworthy. Expert annotators face a different bottleneck: repeatedly verifying and placing many landmarks across large radiograph sets. Automated predictions reduce this manual burden by directing attention to approximate landmark locations, allowing experienced users to focus cognitive effort on fine-tuning placement — reducing total processing time from 10m32s to 6m35s across 10 images. Extrapolated to a representative field season of 1000 radiographs (Stroud *et al*. 2023, 2024), this translates from approximately 15.8 hours of manual annotation to 9.9 hours with LizardMorph — a saving of over 6.5 hours that could determine whether morphometric analyses are feasible within project timelines and budgets.

### 5.4 Generalizability Across Biological Systems

A key design objective was to ensure LizardMorph‘s framework extends beyond lizard radiographs. We demonstrated this by training an ML-Morph model on the *Drosophila* wing dataset from Porto & Voje (2020) — 280 microscopy images with 12 landmarks at wing vein intersections — and deploying it through the LizardMorph interface without modification, confirming that the architecture is agnostic to biological system, imaging modality, and landmark configuration.

Generalizability rests on two requirements: specimens must be imaged in a reasonably consistent position so that the shape predictor can learn reliable spatial patterns, and sufficient manually annotated training images must be available (670 images for the 34-landmark lizard model; 224 for the 12-landmark *Drosophila* model). Systems meeting these requirements span a wide range of biological contexts, including museum specimen photography, histological sections, and veterinary radiography. As specimen imaging becomes increasingly standardized through digitization initiatives such as iDigBio (Nelson & Ellis 2019), the pool of datasets suitable for LizardMorph-style automation will continue to grow.

LizardMorph occupies a distinct niche within the broader ecosystem of morphometric tools. Deep learning frameworks such as DeepLabCut (Mathis *et al*. 2018) and SLEAP (Pereira *et al*. 2022) offer powerful pose estimation but are designed primarily for video analysis and require substantial computational expertise. ML-Morph (Porto & Voje 2020) provides excellent landmark accuracy but lacks a graphical interface for non-programmers. General annotation platforms such as Labelbox and CVAT are not tailored to morphometric workflows or compatible with standard output formats. LizardMorph bridges these gaps by combining a domain-specific ML backend, an interactive correction interface, and direct export compatibility with standard geometric morphometrics software such as MorphoJ (Klingenberg 2011) and geomorph (Adams & Otárola-Castillo 2013), making it a practical tool for biologists collecting morphometric data that lack computational expertise.

### 5.5 Limitations

Several limitations of this study should be considered. First, the performance evaluation involved only two users annotating 10 images each. Although the within-subjects design controls for individual differences and the results were consistent across images, a broader evaluation with more participants spanning a wider range of expertise levels and biological backgrounds.

Second, model accuracy was reduced for spatially inconsistent landmarks — digit tips and scale bar endpoints — representing a genuine limitation for studies in which these features are of primary interest. While human correction mitigates this in practice, more flexible architectures such as convolutional neural networks with spatial attention mechanisms may better handle positional variability, though potentially at the cost of greater training data requirements and computational complexity.

Third, LizardMorph‘s reliance on standardized specimen positioning limits applicability to workflows lacking consistent orientation protocols. Imaging systems with substantial positional variability—such as field photographs of free-ranging animals or unstandardized museum specimens—would fall outside the current pipelinès scope.

Fourth, the current implementation accepts only JPEG images and depends on server-hosted deployment, introducing dependencies on network connectivity and institutional computing infrastructure. Expanding format support and developing lightweight local deployment options would improve accessibility for researchers in resource-limited settings or those working with data that cannot be transmitted to external servers.

### 5.6 Future Directions

Several avenues for future development emerge from this work. Most immediately, we plan to enable researchers to upload their own annotated datasets for custom model training directly through the platform, substantially expanding the range of biological systems LizardMorph can serve. In reference to technical development, deep learning architectures—such as convolutional neural networks or transformer-based pose estimation models—could improve performance on spatially variable landmarks, while active learning strategies could optimize the trade-off between annotation effort and accuracy by prioritizing uncertain predictions for human review and reducing total training data requirements.

Broader adoption would benefit from systematic usability evaluation across multiple laboratories and biological systems, including System Usability Scale assessments and longitudinal studies tracking real-world workflow integration. Such evaluations would provide evidence-based guidance for interface refinements and identify adoption barriers not apparent from small-scale testing. Integration with data standards in biodiversity informatics repositories (e.g., GBIF; ref) would further embed LizardMorph within the digital infrastructure supporting large-scale ecological and evolutionary research.

### 5.7 Conclusion

Manual landmarking remains one of the most persistent methodological bottlenecks in morphological research, constraining the scale and pace of studies across ecology and evolutionary biology. LizardMorph demonstrates that this bottleneck can be substantially alleviated by integrating machine learning automation with human-centered interface design — meeting the dual goals of efficiency and scientific rigor that fully automated or fully manual approaches cannot achieve alone. The framework is intentionally replicable: the same architecture can be adapted to any biological system in which specimens are consistently imaged, without modification to the underlying interface. As ecological and evolutionary research increasingly demands large-scale phenotypic data to address questions about adaptation, phenotypic plasticity, and community assembly, tools that democratize access to high-quality morphometric analysis become essential research infrastructure (Lürig *et al*. 2021). LizardMorph provides a practical template for how ML-assisted annotation tools can expand what is feasible in biological research while preserving the human expertise on which data quality ultimately depends.

## Acknowledgements

This work was supported by the David and Lucille Packard Foundation and the Maxwell/Hanrahan Foundation (both to J.T.S.). This research was supported in part through research cyberinfrastructure resources and services provided by the Partnership for an Advanced Computing Environment (PACE) at the Georgia Institute of Technology, Atlanta, Georgia, USA (RRID:SCR_027619). We also thank the Georgia Tech College of Computing for hosting and supporting the web application resources used in this study.

## Notes

https://github.com/Human-Augment-Analytics/LizardMorph

### Competing Interest Statement

The authors have declared no competing interest.

